# A single-dose of intranasal vaccination with a live-attenuated SARS-CoV-2 vaccine candidate promotes protective mucosal and systemic immunity

**DOI:** 10.1101/2023.04.17.537235

**Authors:** Awadalkareem Adam, Birte Kalveram, John Yun-Chung Chen, Jason Yeung, Leslie Rodriguez, Ankita Singh, Pei-Yong Shi, Xuping Xie, Tian Wang

## Abstract

An attenuated SARS-CoV-2 virus with modified viral transcriptional regulatory sequences and deletion of open-reading frames 3, 6, 7 and 8 (∆3678) was previously reported to protect hamsters from SARS-CoV-2 infection and transmission. Here we report that a single-dose intranasal vaccination of ∆3678 protects K18-hACE2 mice from wild-type or variant SARS-CoV-2 challenge. Compared with wild-type virus infection, the ∆3678 vaccination induces equivalent or higher levels of lung and systemic T cell, B cell, IgA, and IgG responses. The results suggest ∆3678 as an attractive mucosal vaccine candidate to boost pulmonary immunity against SARS-CoV-2.

## Main Text

Severe acute respiratory syndrome coronavirus 2 (SARS-CoV-2), the cause of the coronavirus disease 2019 (COVID-19) pandemic, has resulted in more than one million deaths in the United States over the past three years. In response to the pandemic, there has been rapid development of SARS-CoV-2 vaccines. Currently, four vaccines have been granted emergency use authorization (EUA) by the FDA. Although the vaccines are highly effective against severe disease, their efficiency has been challenged by the increasing rates of variants of concern (VOCs) that are characterized by increased viral transmissibility and immune evasion [1]. Continuous work is needed to optimize existing vaccine platforms and to develop more effective novel vaccines.

Parenteral injection route is currently used for the delivery of approved COVID-19 vaccines. Such administration route triggers limited local respiratory tract immune responses, particularly against VOCs [2]. Compared to the parenteral route, delivery of antigens to the sites of infection induces mucosal immune responses in the respiratory tract, including IgA and resident memory B and T cells, which provide additional layers of protection [3]. Thus, intranasal immunization, which leads to the induction of antigen-specific immunity in both mucosal and systemic immune compartments [4], would be more effective in controlling viral infection and disease [5, 6]. Here we test this hypothesis by mucosal immunization of K18-hACE2 mice with a highly attenuated SARS-CoV-2 containing modified viral transcriptional regulatory sequences and deletion of open-reading frames 3, 6, 7 and 8 (∆3678). This vaccine candidate was previously shown to protect hamsters from SARS-CoV-2 infection and transmission after a single-dose vaccination [7]. Our current study shows that a single-dose intranasal vaccination of K18-hACE2 mice with

∆3678 induces potent mucosal and systemic T cell and humoral immune responses at similar or higher levels than WT virus infection, leading to a full protection of all immunized mice from WT and VOC SARS-CoV-2 challenge.

K18-hACE2 mice were immunized intranasally (i.n.) with 2×10^3^ PFU of ∆3678 virus. Mice vaccinated with the wild-type (WT) SARS-CoV-2 USA-WA1/2020 or PBS (mock) were used as controls (**Fig. 1A**). All animals vaccinated with the ∆3678 virus or PBS survived 28 days post-vaccination (DPV) and displayed neither weight loss nor clinical signs (**Fig. 1B**); whereas 7.6% of the animals infected with the same dose of WT virus succumbed to disease between 7 and 12 DPV (**data not shown**). About one-third of the mice inoculated with the WT virus showed weight loss after day 7 (**Fig. 1B**). At 28 DPV, blood, bronchoalveolar lavage (BAL) fluid, lung, and spleen samples were collected to determine the virus-induced immune responses. In the lung, both WT and ∆3678 virus-inoculated mice displayed a Th1-prone immune response.

**Figure 1.**
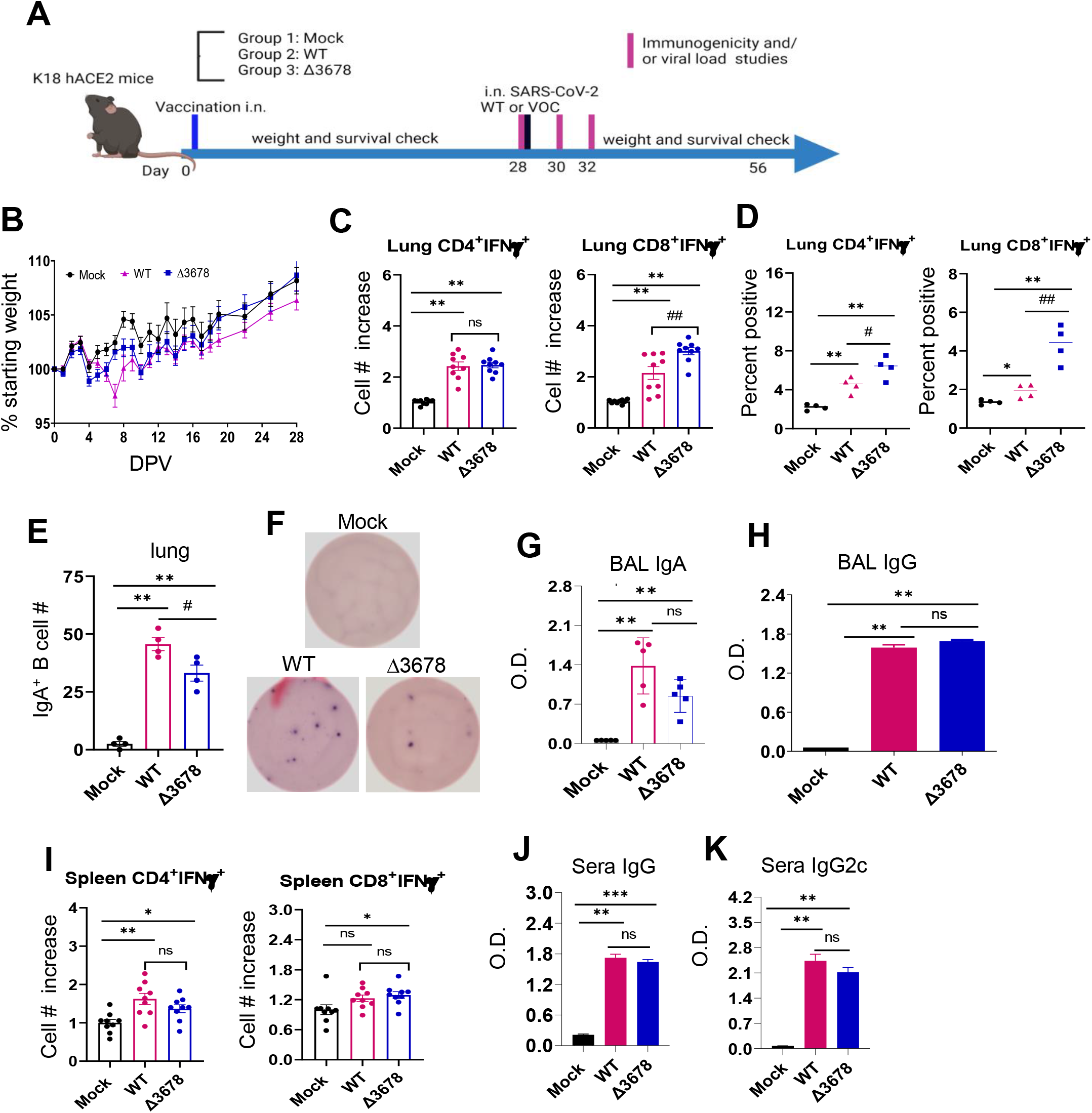
∆3678 SARS-CoV-2 induced strong mucosal and systemic immune responses in K18-hACE2 mice one month post intranasal vaccination. **A**. Study design and vaccination timeline. **B**. Weight loss is indicated by percentage using the weight on the day of vaccination as 100%. **C-D & I**. Lung leukocytes (**C-D**) or splenocytes (**I**) were cultured *ex vivo* with S peptide pools for 6 h, and stained for IFN-γ, CD3, and CD4 or CD8. Fold increase of IFN-γ^+^ CD4^+^ and CD8^+^ T cells expansion compared to the mock vaccinated group is shown (**C** & **I**). **D**. Percent positive of IFN-γ^+^ CD4^+^ and CD8^+^ T cells among lung T cells is shown. **E-F**. Lung leukocytes were stimulated *in vitro* for 7 days with immunostimulatory agents (R848 plus rIL-2) and seeded onto ELISPOT plates coated with SARS-CoV-2 RBD. Frequencies of SARS-CoV-2 RBD specific IgA-secreting lung B cells per 10^6^ input cells in MBC cultures. **G-H**. IgA (**G**) and IgG (**H**) titers in BAL presented as O.D. values by ELISA. **J-K**. Sera IgG (**J**) and IgG2C (**K**) titers presented as O.D. values by ELISA. ** *P* < 0.01, or \**P* < 0.05 compared to mock group. ^##^*P* < 0.01, or ^#^*P* < 0.05 compared to WT group.

Compared to the WT virus group, CD4^+^ T cells from the ∆3678-vaccinateded mice had similar or higher levels of IFNγ^+^ production, as indicated by the percentage and total cell number increase; whereas CD8^+^IFNγ^+^ T cells displayed an increase in both percentage and total cell number (**Fig. 1C, D**). Both WT and ∆3678 inoculations triggered high RBD-specific IgA^+^ B cell responses, though the latter group had 25% lower response (**Fig. 1E, F**). Furthermore, comparable levels of SARS-CoV-2-specific IgA or IgG antibodies were detected in the BAL fluid of the WT and ∆3678 groups (**Fig. 1G, H**). In the periphery of WT- and ∆3678-vaccinated mice, Th1-prone immune responses were also noted in the spleen. CD4^+^IFNγ^+^ T cells in the ∆3678-vaccinated mice had similar cell number as the WT virus group with a decrease on the percentage, whereas CD8^+^ T cells of the ∆3678 group produced IFN-γlevels similar to the WT virus by total cell number and percentage (**Fig. 1I**, and **Supplementary Fig.1A)**. Additionally, both WT and ∆3678 viruses induced strong SARS-CoV-2-specific IgG and Th1-prone IgG2c in the sera; no differences in antibody titers were observed between the two groups (**Fig. 1J, K**).

Overall, the results suggest that although ∆3678 was highly attenuated in K18-hACE2 mice, it induces potent mucosal and systemic T cell and humoral immune responses at equivalent or higher levels than WT virus.

The protective efficacy of the ∆3678 mutant was determined by challenging the vaccinated mice with 10^4^ PFU of the WT USA-WA1/2020 virus on 28 DPV. Animals that had previously been inoculated with the WT or ∆3678 virus survived the challenge, while 4 out of 9 of mock-vaccinated animals succumbed to disease between days 7 and 8 post-challenge (**Fig. 2A**). Mice vaccinated with either virus displayed no weight loss after the challenge, while the naïve animals lost up to 20% of body weight by day 7 post-challenge (**Fig. 2B**). Mice that had previously been inoculated with WT or ∆3678 virus had no detectable virus in their lungs or tracheae on days 2 and 4 and significantly diminished viral loads in nasal washes at day 2 post-challenge (**Fig. 2C-E**). To determine protective efficacy against more recently circulating variants, we also performed a challenge with Omicron BA.5 virus – a VOC with low susceptibility to antibody neutralization when tested against previous variant infected convalescent sera or vaccinated sera [8]. Strikingly, none of the mice that had been inoculated with the ∆3678 mutant had detectable virus in the lung or trachea on day 2 post-challenge while the naïve animals had viral loads of over 10^6^ PFU/g lung tissue (**Fig. 2F**). Thus, vaccination with the ∆3678 mutant has equivalent protective efficacy as WT virus against subsequent WT virus or VOC challenge in mice.

**Figure 2.**
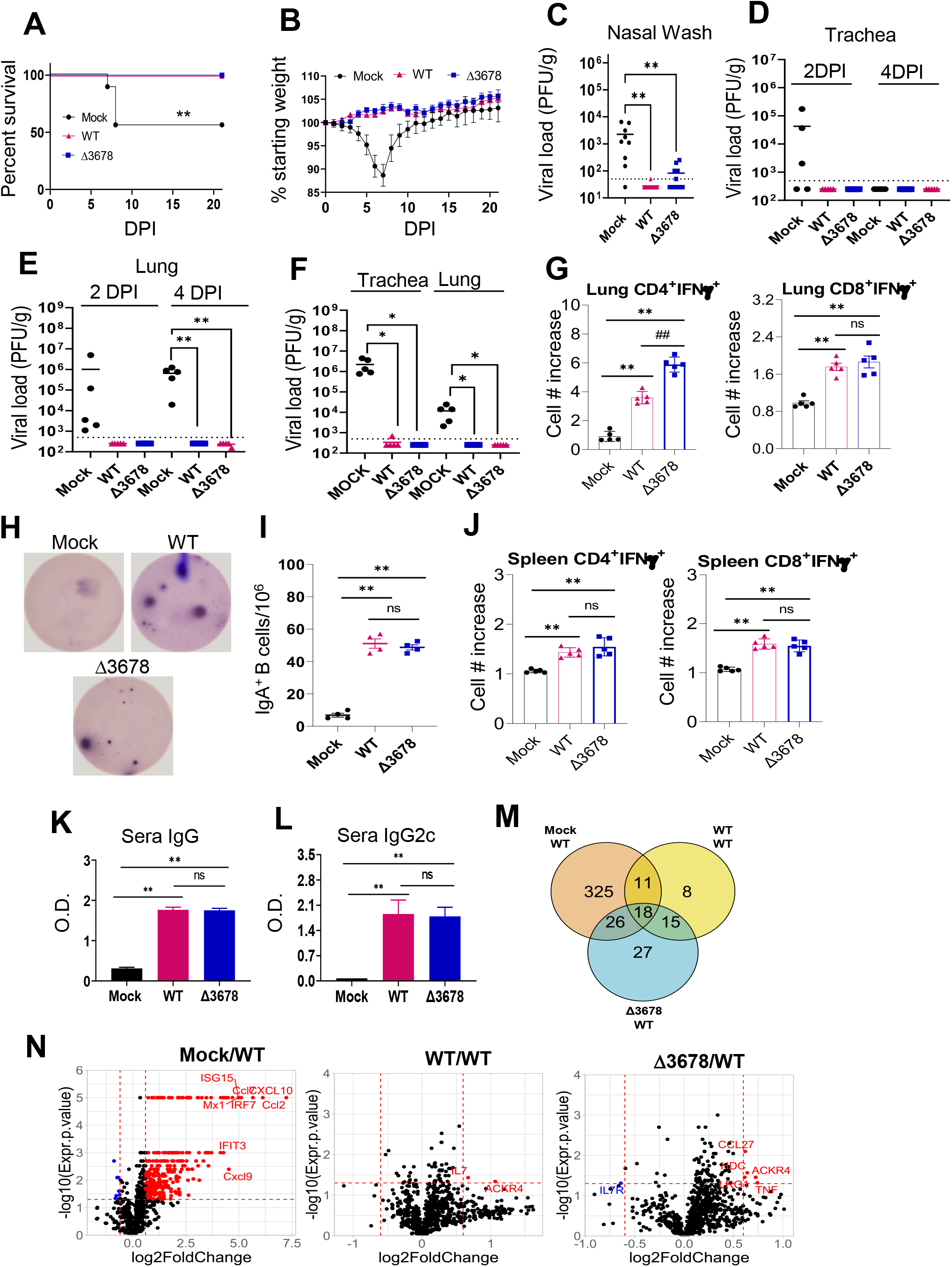
Intranasal vaccination of ∆3678 SARS-CoV-2 protects mice from WT or VOC virus challenge via induction of lung resident and systemic T cell and humoral immune responses. K18-hACE2 mice were intranasally immunized with mock (PBS), WT USA-WA1/2020 SARS-CoV-2, or ∆3678 virus. At 28 DPV, mice were intranasally challenged with 10^4^ PFU of WT USA-WA1/2020 virus. Mice were monitored daily for survival (**A**) and weight loss (**B**). Weight loss is indicated by percentage using the weight on the day of infection as 100%. (**C-E**) SARS-CoV-2 viral loads in nasal washes, trachea, and lungs were measured by plaque assays at indicated days post-infection (DPI). **F**. Viral loads in trachea and lung at day 2 post-challenge with Omicron BA.5 SARS-CoV-2. **G & J**. Lung leukocytes (**G**) or splenocytes (**J**) were cultured *ex vivo* with spike peptide pools for 5 h, and stained for IFN-γ, CD3, and CD4 or CD8. Fold increase of IFN-γ^+^ CD4^+^ and CD8^+^ T cells expansion compared to the mock-vaccinated group is shown. **H-I**. Lung leukocytes were stimulated *in vitro* for 7 days with immunostimulatory agents (R848 plus rIL-2) and seeded onto ELISPOT plates coated with SARS-CoV-2 RBD. Frequencies of SARS-CoV-2 RBD-specific IgA-secreting lung B cells per 10^6^ input cells in MBC cultures are shown. K**-L**. Sera IgG (**K**) and IgG2C (**L**) titers presented as O.D. values by ELISA. **M-N**. NanoString data for mouse lungs (harvested on day 4 post challenge) from designated vaccination groups. All comparisons are between a challenged group and unvaccinated/unchallenged (mock/mock) controls. **M**. Venn diagram of differentially expressed genes (unadjusted *p* value ≥0.05) between the three vaccination conditions. **N**. Volcano plots. The horizontal dotted line corresponds to an unadjusted *p* value cutoff of 0.05 and vertical lines correspond to −0.6 and 0.6 log2 (fold change). ** *P* < 0.01, or \**P* < 0.05 compared to mock group. ^##^*P* < 0.01, or ^#^*P* < 0.05 compared to WT group.

To further assess the ∆3678-induced protective immunity, blood, lung and spleen samples were collected on day 4 post-challenge with the WT virus. In the lung, CD4^+^ IFN-γ^+^ T cells of the ∆3678 group had about 50% increases in total cell number and the percentage compared to the WT virus-vaccinated mice; while the number and percentage of CD8^+^ IFN-γ^+^ T cells were similar between the two groups. Compared to the mock group, the number of CD4^+^IFN-γ^+^ and CD8^+^ IFN-γ^+^ T cells increased 550% and 89% respectively (**Fig. 2G** and **Supplementary Fig. 1B**).

Furthermore, ∆3678-vaccinated mice produced a similar number of IgA^+^-expressing B cells as the WT virus-vaccinated group in response to virus challenge (**Fig. 2H-I**). In the periphery, splenic CD4^+^ and CD8^+^ T cells of both the WT- and ∆3678-vaccinated mice induced more IFN-γproduction than the mock group at day 4 post-challenge but no differences in IFN-γlevels were detected between the WT- and ∆3678-vaccinated mice (**Fig. 2J** and **Supplementary Fig. 1C)**. Lastly, strong but similar titers of SARS-CoV-2-specific IgG and IgG2c antibodies were detected in the sera of both the WT- and ∆3678-vaccinated groups (**Fig. 3K-L**).

We next investigated the underlying immune mechanisms of ∆3678-induced host protection. By using the probe-based nCounter Analysis system, we compared the mRNA expression of 785 host genes from lung homogenates of mock-, WT-, and ∆3678-vaccinated mice at day 4 post-challenge with the WT virus. The three groups of lung homogenates were labeled mock/WT, WT/WT, and ∆3678/ WT, respectively. All samples were compared with lung samples collected from the unvaccinated and unchallenged group (mock/mock). Twenty canonical pathways (**Supplementary Fig. 2**) were identified from comparison analysis by the Ingenuity Pathway Analysis (IPA). Among them, pathogen-induced cytokine storm signaling pathways, pyroptosis signaling pathway, necroptosis signaling pathway, and type I diabetes signaling, which are associated with the severity of SARS-CoV-2 infection [9–11], were induced in the mock/WT group but diminished in the WT/WT and ∆3678/WT groups, suggesting that both WT and ∆3678 viruses induced host protection against WT virus challenge. Furthermore, 15 overlapping differentially expressed genes (DEGs) were observed in both WT/WT and ∆3678/WT groups (**Fig. 2M, N**, and **Table 1**). The DEGs include (i) genes associated with lung tissue resident memory T cells (Trm) development and maintenance (e.g., *Il7* and *Tgfb3)* [12], (ii) genes associated with CD8 Trm-initiated activities in the lung (e.g., *Gzmk, Fcrgrt*) [13], and (iii) genes associated with T cell recruitment into the lung (e.g., *Ackr4*) [14]. These results further support that mucosal vaccination with the ∆3678 mutant protects mice from the SARS-CoV-2 challenge by promoting lung resident memory T cell responses.

**Table 1.** Overlapping differentially expressed genes with fold change and p values for each group.

| Gene | WT/WT | | $\Delta 3678$/WT | |
| --- | --- | --- | --- | --- |
|  | Fold Change | P Value | Fold Change | P Value |
| *Ackr4* | 2.093 | 0.046 | 1.652 | 0.035 |
| *Anpep* | 1.198 | 0.024 | 1.300 | 0.017 |
| *Atf4* | -1.399 | 0.008 | -1.220 | 0.047 |
| *Ccl6* | 1.267 | 0.033 | 1.341 | 0.017 |
| *Ctsv* | 1.131 | 0.050 | 1.177 | 0.022 |
| *Fcgrt* | 1.314 | 0.022 | 1.333 | 0.024 |
| *Gzmk* | 1.332 | 0.022 | 1.384 | 0.049 |
| *Il7* | 1.593 | 0.037 | 1.363 | 0.019 |
| *Neo1* | 1.137 | 0.045 | 1.248 | 0.009 |
| *Neu1* | 1.137 | 0.017 | 1.115 | 0.043 |
| *Os9* | 1.108 | 0.020 | 1.138 | 0.004 |
| *Ptpn4* | 1.142 | 0.016 | 1.200 | 0.005 |
| *Tgfb3* | 1.072 | 0.018 | 1.180 | 0.042 |
| *Timp2* | 1.084 | 0.034 | 1.162 | 0.002 |
| *Ywhaq* | 1.102 | 0.007 | 1.107 | 0.002 |

NanoString data for mouse whole lung day 4 from designated vaccination groups. All comparisons are between a challenged group and unvaccinated/unchallenged (mock/mock) controls.

In summary, our results consistently indicate that ∆3678 SARS-CoV-2 is highly attenuated in K18-hACE2 mice. A single-dose of intranasal vaccination with this vaccine candidate protects from WT and VOC virus challenge via induction of SARS-CoV-2-specific mucosal and systemic cell-mediated and humoral immune responses. We also demonstrated that mucosal delivery of the live-attenuated vaccine candidate promotes the development and functional activation of pulmonary Trms. Therefore, the ∆3678 virus can serve as an attractive candidate for future vaccination via the mucosal route either by itself or by a prime-pull strategy, leading to enhanced and/or durable mucosal immunity in the respiratory tracts [15, 16].

## Materials and Methods

### Viruses and Cells

African green monkey kidney epithelial Vero-E6 cells (laboratory-passaged derivatives from ATCC CRL-1586) were grown in Dulbecco’s modified Eagle’smedium (DMEM; Gibco/Thermo Fisher) with 10% fetal bovine serum (FBS, HyClone Laboratories) and 1% antibiotic/streptomycin (P/S, Gibco). Vero-E6-TMPRSS2 cells were purchased from SEKISUI XenoTech, LLC, and maintained in 10% fetal bovine serum (FBS; HyClone Laboratories), and 1% P/S and 1 mg/ml G418 (Gibco). All cells were maintained at 37°C with 5% CO_2_. The infectious clones derived USA-WA1/2020 SARS-CoV-2, ∆3678, and the BA.5 variant were generated as previously described [7, 17, 18].

### Animal Studies

Female heterozygous K18-hACE2 C57BL/6J mice (strain: B6.Cg-Tg(K18-ACE2)2Prlmn/J) were obtained from The Jackson Laboratory and were infected intranasally (i.n.) with 2×10^3^ PFU of either WT WA1 or ∆3678 infectious clone-derived viruses or were mock-infected with DPBS. Mice were between 8 and 10 weeks old at the time of initial infection and were age-matched within experimental cohorts. Intranasal challenge with 1×10^4^ PFU of WT WA1 or BA.5 virus was performed on day 28 after the initial infection. For each infection animals were anesthetized with isoflurane (Piramal). Bronchoalveolar lavage (BAL) fluid, nasal washes, whole lungs, and whole spleens were collected from one subset of mice for immunological analysis. From a different subset of mice, the trachea and cranial right lobe were collected in DPBS for enumeration of viral loads via plaque assay, and the middle and caudal lobes were collected in TRIzol (Invitrogen) for RNA extraction.

### Ethic Statement

All animal handling was performed under animal biosafety level 3 (ABSL3) conditions at the Galveston National Laboratory and in accordance with guidelines set by the Institutional Animal Care and Use Committee of the University of Texas Medical Branch.

### Plaque assay

Collected lung lobes were homogenized in 1 ml of PBS at 6,000 rpm for 60 s using a Roche MagNA Lyser instrument before titration. The lung homogenates were clarified by centrifugation at 12,000 rpm for 3 min. The supernatants were collected for titration using a standard plaque assay. Briefly, approximately 1.2×10^6^ Vero-E6-TMPRSS2 cells were seeded to each well of 6-well plates and cultured at 37 °C, 5% CO2 for 16 h. The virus was serially diluted in DMEM with 2% FBS and 200 μl diluted viruses were transferred onto the monolayers. The viruses were incubated with the cells at 37 °C with 5% CO_2_ for 1 h. After the incubation, an overlay medium was added to the infected cells per well. The overlay medium contained MEM with 2% FBS, 1% penicillin/streptomycin, and 1% Sea-plaque agarose (Lonza, Walkersville, MD). After a 2.5-day incubation, the plates were stained with neutral red (Sigma-Aldrich) and plaques were counted on a lightbox.

### Antibody ELISA

ELISA plates were coated with 100 ng/well recombinant SARS-CoV-2 RBD protein (RayBiotech) overnight at 4°C. The plates were washed twice with PBS, containing 0.05% Tween-20 (PBS-T) and then blocked with 8% FBS for 1.5 h. Sera or BAL samples were diluted 1:100 or undiluted in blocking buffer and were added for 1 h at 37°C. Plates were washed 5 times with PBS-T. Goat anti-mouse IgG (Sigma, MO, USA), or goat anti-mouse IgG2C (Southern Biotech) conjugated with horseradish peroxidase (HRP) or alkaline phosphatase was added at a 1:2,000 dilutions for 1 h at 37ºC followed by adding TMB (3, 3, 5, 5′-tetramethylbenzidine) peroxidase substrate (Thermo Scientific) or *p*-nitrophenyl phosphate (Sigma-Aldrich), and the intensity was read at an absorbance of 450 or 405 nm. To determine IgA titer, HRP-conjugated goat anti-mouse IgA (Southern Biotech) was added as the secondary antibody at a 1:2,000 dilution for 1 h at 37ºC, followed by adding TMB peroxidase substrate (Thermo Scientific) for about 15 min. The reactions were stopped by 1M sulfuric acid, and the intensity was read at an absorbance of 450 nm.

### B cell ELISPOT

ELISPOT assays were performed as previously described [19]. Briefly, Millipore ELISPOT plates (Millipore Ltd, Darmstadt, Germany) were coated with 100 µl rSARS-CoV-2 spike protein (R&D Systems). To detect total IgA^+^-expressing B cells, the wells were coated with 100 µl of anti-mouse IgA capture Ab (Mabtech In). Cells were added in duplicates to assess total IgA ASCs or SARS-CoV-2 specific B cells. The plates were incubated overnight at 37°C, followed by incubation with biotin-conjugated anti-mouse IgA (Mabtech In) for 2 h at room temperature, then 100 µL/well streptavidin-ALP was added for 1 h. Plates were developed with BCIP/NBT-Plus substrate until distinct spots emerge, washed with tap water, and scanned using an ImmunoSpot 6.0 analyzer and analyzed by ImmunoSpot software (Cellular Technology Ltd).

### Intracellular cytokine staining (ICS)

Splenocytes or lung leukocytes were incubated with SARS-CoV-2 spike peptide pools (1μg/ml, Miltenyi Biotec) for 6 h in the presence of GolgiPlug (BD Bioscience). Cells were stained with antibodies for CD3, CD4, or CD8, fixed in 2% paraformaldehyde, and permeabilized with 0.5% saponin before adding anti-IFN-γ(Thermo Fisher Scientific). Samples were processed with a C6 Flow Cytometer instrument. Dead cells were excluded based on forward and side light scatter. Data were analyzed with a CFlow Plus Flow Cytometer (BD Biosciences).

### Analysis of nCounter Analysis System (NanoString) Data

Lung samples were homogenized in Trizol (Thermo Fisher Scientific). Total RNA was purified using the Direct-zol-96 MagBead RNA kit (Zymo) with a KingFisher Apex System (Thermo Fisher Scientific). RNA samples were normalized to 20 ng/μL and followed by analysis using the nCounter Pro Analysis System and the nSolver Analysis Software. Plots were made using R version 4.1.2. An un-adjusted p-value cutoff of 0.05 was used to determine differentially expressed genes (DEGs) for the three conditions. An additional log2 fold change cutoff of + or – 0.6 was used to limit labeling of up-and down-regulated genes in volcano plots although the Mock/WT condition had too many\ genes to label. The ‘ggvenn’ package version 0.1.9 was used to create Fig. 2M. Ingenuity pathway analysis (version 84978992) core analysis was performed on data (gene name, un-adjusted p-value, and fold change) from each condition, evaluating based on expression fold changes. The option “User Data Set” was used as the reference set. Data on canonical pathways for Supplementary Fig. 2 were derived from a comparison analysis. −log(p-values) and z-scores were used to generate bubble plots, using ascending p-value or descending −log(p-value) for ordering.

### Statistical analysis

Values for viral load, cytokine production, antibody titers, B and T cell response experiments were compared using Prism software (GraphPad) statistical analysis and were presented as means ± SEM. *P* values of these experiments were calculated with a non-paired Student’s t test.

## Supporting information

AdamKalveram Supplementary Figures

## DATA AVAILABILITY

All data generated or analyzed during this study are included in this published article (and its supplementary information files).

## ACKNOWLEDGEMENTS

This work was supported in part by NIH grants R01AI127744 (T.W.), and R01 NS125778 (T.W.). P.Y.S. was supported by NIH contract HHSN272201600013C and awards from the Sealy & Smith Foundation and the Kleberg Foundation.

## AUTHOR CONTRIBUTIONS

A.A., B.K., J.C., L.R. A.S., X.X., performed the experiments. P.Y.S., X.X., and T.W., designed the experiment. A.A., B.K., J.Y., X.X., and T.W. analyzed the data. T.W. wrote the initial draft of the manuscript and other coauthors provided editorial comments.

## SUPPLEMENTARY FIGURE LEGENDS

**Supplementary Figure 1. SARS-CoV-2 ∆3678 mutant induced strong mucosal and systemic T cell responses in K18-hACE2 mice one month post intranasal vaccination followed by WT virus challenge**. Lung leukocytes (**B**) or splenocytes (**A & C**) collected at 28 DPV (**A**) or day 4 post-challenge (**B-C**) were cultured *ex vivo* with S peptide pools for 5 h, and stained for IFN-γ, CD3, and CD4 or CD8. Percent positive of IFN-γ^+^ CD4^+^ and CD8^+^ T cells among lung or spleen T cells is shown. ** *P* < 0.01, or \**P* < 0.05 compared to mock group. ^##^*P* < 0.01, or ^#^*P* < 0.05 compared to WT group.

**Supplementary Figure 2. NanoString data for mouse whole lung day 4 post-challenge in the vaccinated mice**. All comparisons are between a challenged group and unvaccinated/unchallenged (mock/mock) controls. Bubble plot of 20 canonical pathways ordered by ascending p value from comparison analysis in Ingenuity Pathway Analysis. Dot size corresponds to –log(p value) where larger dots indicate more significant p values. Color intensity corresponds to activation z-score and all pathways are either activated or demonstrate no change. Gray indicates values that could not be calculated.

