## Supplementary figures and images for "A single-dose of intranasal vaccination with a live-attenuated SARS-CoV-2 vaccine candidate promotes protective mucosal and systemic immunity"

### AdamKalveram Supplementary Figures

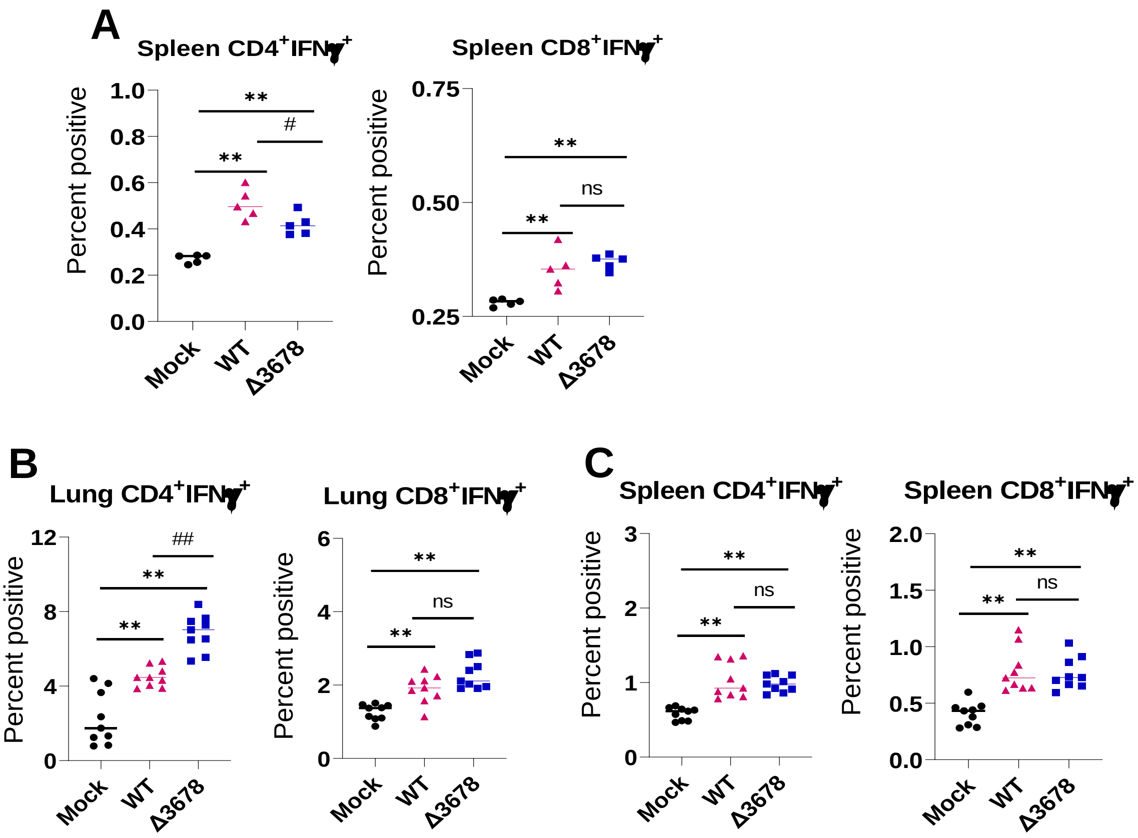


Suppl Figure 1

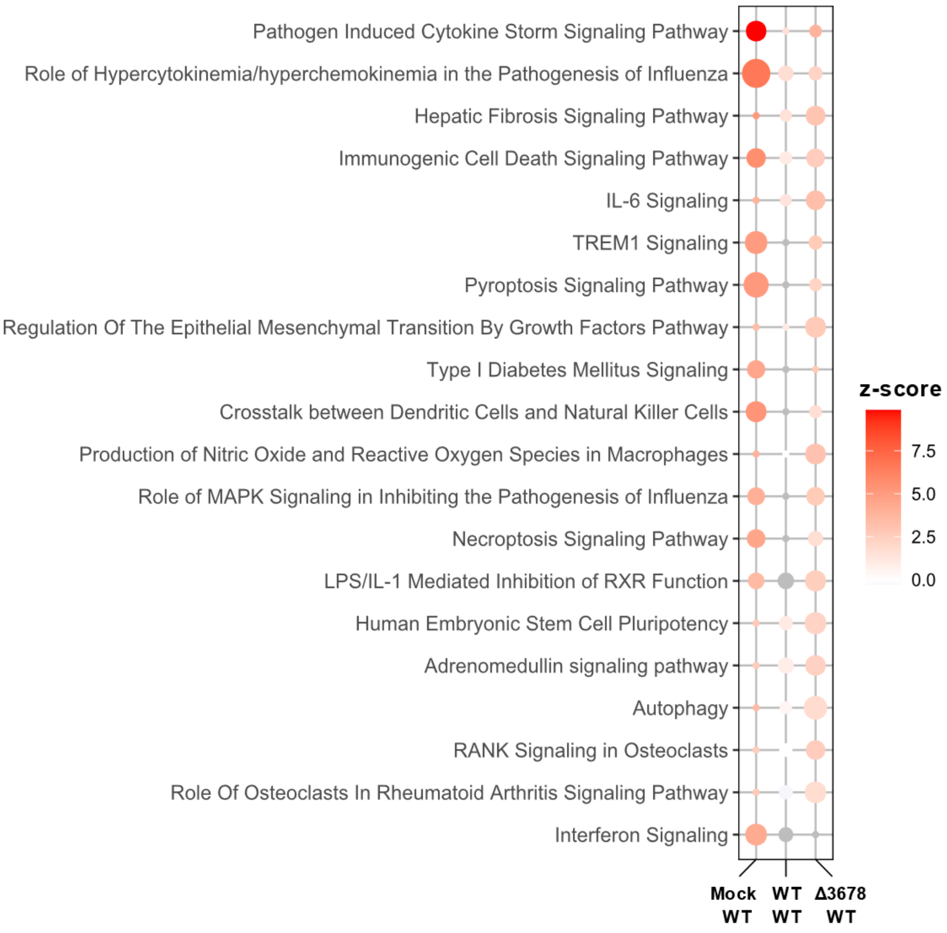


**Suppl Figure 2**
